# EXTRACT 2.0: text-mining-assisted interactive annotation of biomedical named entities and ontology terms

**DOI:** 10.1101/111088

**Authors:** Evangelos Pafilis, Rūdolfs Bērziņš, Lars Juhl Jensen

## 1 INTRODUCTION

Databases increasingly rely on text-mining tools to support the curation process. The BioCreative interactive annotation task recently evaluated several such tools and found our tool EXTRACT to perform favorably in terms of usability and accelerated curation by 15–25% (Wang *et al,* 2016).

The original version of EXTRACT was designed to support annotation of metagenomic samples with semantically controlled environmental descriptors (Pafilis *et al.,* 2016). For this reason, it focused on named entity recognition of terms from the Environment Ontology (ENVO) (Buttigieg *et al.,* 2016) and ontologies relevant for describing host organisms (https://www.ncbi.nlm.nih.gov/taxonomy), tissues (Placzek *et al,* 2017), and disease states (Kibbe *et al.,* 2015).

## 2 EXPANDED SCOPE OF THE TOOL

EXTRACT 2.0 expands the scope of the tool in several new directions with the aim to make it more broadly useful.

We expanded the scope from covering only diseases to covering phenotypes in general. To this end, we complemented the existing disease dictionary with terms from the Mammalian Phenotype Ontology (MPO) (Smith and Eppig, 2012). To avoid redundancy in the dictionary, we excluded MPO terms that clashed with terms already in the disease dictionary. To improve recall, we added plural and adjective endings to the names and generated variants of the form *pronoun of noun* from names of the form *noun pronoun*.

To cover also important concepts of molecular and cellular biology, we further expanded the dictionary with Gene Ontology (GO). The names from GO were processed similar to those from MPO to generate variants and improve recall.

In addition to adding more biomedical ontologies, we have expanded the tool with named entity recognition of molecular entities. To this end, we have included dictionaries of protein-coding and non-coding RNA (ncRNA) genes from STRING (Szkararczyk *et al.,* 2017) and RAIN (Junge *et al.,* 2017), respectively. We have furthermore added a dictionary of drugs and other small molecule compounds from the STITCH database (Szklarczyk *et al.,* 2016).

Together, these additional types of entities have made EXTRACT 2.0 potentially useful for many more tasks than just annotation of metagenomic samples. For example, it can be used to help annotate both proteins and ncRNAs with functions, processes, subcellular localization, tissue expression, and associated diseases. The tool, API, and documentation are freely accessible at http://extract.jensenlab.org.

